# Disrupted Top-Down Modulation as a Mechanism of Impaired Multisensory Processing in Autistic Children

**DOI:** 10.1101/2025.11.13.688243

**Authors:** Theo Vanneau, John J. Foxe, Shlomit Beker, Sophie Molholm

## Abstract

Atypical sensory processing is a core feature of autism, particularly when integration across sensory modalities is required. The neural mechanisms underlying these multisensory differences remain unclear. We recorded high-density EEG while autistic children aged 8–13 (AU; n=40), unaffected siblings of autistic children (SIB; n=26), and non-autistic controls (NA; n=36) performed a simple reaction-time task to auditory (A), visual (V), and audiovisual (AV) stimuli. Analyses targeted event-related potentials (ERPs; P1/N1/P2), alpha-band event-related desynchronization (α-ERD), and long-range theta-band functional connectivity (weighted phase-lag index, wPLI). Across all unisensory measures (ERPs, α-ERD, and connectivity), groups did not differ, indicating broadly comparable unisensory processing. By contrast, multisensory integration (MSI; operationalized for ERPs and α-ERD as AV − (A+V)) differed across groups: NA children showed significant ERP MSI over parieto-central sites that was absent in AU and SIB; and α-ERD MSI was present in all groups but significantly reduced in AU, with SIB showing an intermediate profile. Connectivity analyses revealed that AV theta-band fronto–parieto-occipital coupling was reduced in autistic relative to non-autistic children, consistent with weaker large-scale coordination during multisensory processing. Together, these results point to a multisensory-specific deficit in autism spanning early sensory encoding, posterior α-ERD, and fronto–posterior coupling. The convergence of results supports a mechanistic account of disrupted multisensory influences on sensory processing due to reduced multisensory attentional orientation. Intermediate SIB profiles suggest inherited liability for these neural phenotypes. These results help explain well-documented behavioral MSI differences in autism by linking impaired early enhancement with attenuated top-down control of sensory cortex.

## Introduction

Our surroundings are filled with myriad sensory cues that carry critical information about the world. Specialized sensory organs transduce the diverse physical energies from these cues into neural signals that are relayed to the brain through sensory-specific pathways to support perception, cognition, and behavior. When sensory inputs from multiple modalities occur together in time and space, they often originate from the same event and convey redundant or complementary information (Stein et al., 1988). Integrating neural signals from these multisensory cues enhances stimulus detection, localization, and identification, allowing the brain to construct a coherent representation of the environment even when individual sensory channels are noisy or degraded (Molholm & Foxe, 2010; Stein, 1998; Stein & Arigbede, 1972). Experimentally, this process manifests as improved behavioral performance and distinct neural responses to multisensory compared to unisensory stimulation that cannot be explained by simple linear summation or winner-takes-all mechanisms (Diederich & Colonius, 2004; Molholm et al., 2002; Ross et al., 2022; Ross et al., 2007; Wallace et al., 1998).

A significant body of research indicates that autistic children^1^ derive less benefit from multisensory inputs than their non-autistic peers. They often show reduced behavioral facilitation and fewer multisensory illusions accompanied by atypical neural modulation during multisensory processing (Brandwein et al., 2013; Ross et al., 2015; Ross et al., 2024; Russo et al., 2010; Stefanou et al., 2024). This diminished integration of concurrent sensory inputs into unified, meaningful percepts may contribute to the frequent experience of sensory overload in autism (Leekam et al., 2006; Schaaf et al., 2025). Furthermore, reduced benefit from multisensory cues, especially under degraded signal conditions, likely exacerbates communication challenges (Beker et al., 2018; Stevenson et al., 2017; Vanneau, Crosse, et al., 2025), a hallmark of the condition.

Electroencephalographic (EEG) studies reveal that early sensory cortical activity is modulated by multisensory stimulation (Brandwein et al., 2011; Molholm et al., 2002), with neural responses to multisensory stimuli that deviate from the linear sum of their unisensory components (e.g., AV ≠ A + V) providing a signature of multisensory integration (MSI). In autism, however, this MSI effect is frequently reduced or absent (Brandwein et al., 2013; Russo et al., 2010; Stefanou et al., 2024), offering neurophysiological evidence of atypical multisensory processing. Complementing event-related potentials (ERPs), event-related spectral analyses decompose the EEG signal into frequency bands that reveal oscillatory dynamics underlying multisensory processing. In particular, alpha-band activity (7–13 Hz) has been closely linked to sensory processing and attentional control (Foxe & Snyder, 2011; Hanslmayr et al., 2011; Klimesch et al., 2007; Senkowski & Engel, 2024) and modulates in visual cortices in response to multisensory stimulation (Cooke et al., 2019; Romei et al., 2012). A decrease in alpha power following stimulus onset (known as alpha event-related desynchronization: α-ERD) indexes increased cortical excitability and engagement of sensory regions (Hanslmayr et al., 2011; Pfurtscheller & Klimesch, 1992) and is modulated by attention and task demands (Capotosto et al., 2009; Li et al., 2013; Peng et al., 2015; Sauseng et al., 2005; Zhozhikashvili et al., 2022). Recently, Matyjek et al. (Matyjek et al., 2025) reported reduced audiovisual speech–evoked α-ERD in autistic adults, suggesting that alpha modulation may serve as a sensitive marker of altered multisensory influences on sensory processing in autism.

Atypical MSI in autism may arise, at least in part, from disrupted top-down attentional modulation that fails to appropriately regulate sensory cortical activity. Multisensory events engage distributed neural networks that require coordinated information exchange, both among sensory cortices and between these regions and higher-order integrative/attentional systems (Fiebelkorn et al., 2010; Mercier et al., 2015; Molholm et al., 2007; Molholm et al., 2002). Temporally synchronous multisensory signals are particularly effective at capturing attention (Bahrick & Lickliter, 2000), whereas differences in multisensory attention have been linked to both language and symptom severity in autism (Todd & Bahrick, 2023). Theta-band (4–7 Hz) oscillations, especially over frontal and posterior regions, are often enhanced during audiovisual processing and may mediate communication between attentional control and sensory systems (Michail et al., 2022). Within this framework, during multisensory stimulation, fronto-parietal attention and salience networks may exert top-down influences via theta-band (4–7 Hz) synchronization, transiently releasing posterior alpha inhibition (i.e., producing α-ERD) and thereby amplifying early sensory responses (Cavanagh & Frank, 2014; Foxe & Snyder, 2011; Hanslmayr et al., 2011; Keil & Senkowski, 2018; Klimesch et al., 2007; Senkowski et al., 2008). Given consistent evidence for reduced or delayed long-range connectivity in autism (Long et al., 2016; Monk et al., 2009; Muller et al., 2011), particularly between frontal and posterior regions (Chen et al., 2024; Jao Keehn et al., 2021), blunted attentional modulation of parieto-occipital α-ERD may contribute to diminished multisensory enhancement.

Motivated by this framework, we assessed multisensory processing using event-related potentials, parieto-occipital α-ERD, fronto-posterior and temporal-posterior theta connectivity within the same paradigm to examine whether early sensory encoding differences co-vary with local excitability and large-scale network coordination. Leveraging the full dataset from this study, we further asked whether typically developing individuals with an autistic sibling (SIBs), who share familial risk but are clinically unaffected, exhibit intermediate neural profiles consistent with endophenotypic liability. This approach allows us to determine whether neural alterations observed in autism reflect disorder-specific effects or broader familial traits linked to genetic risk.

We hypothesized that, relative to non-autistic controls, autistic children would exhibit reduced multisensory enhancement of early ERPs, attenuated α-ERD modulation, and weakened fronto-posterior theta connectivity, with SIBs showing intermediate patterns. To test these hypotheses, we analyzed high-density EEG data collected while children performed a simple reaction-time task involving randomly interleaved auditory (A), visual (V), and audiovisual (AV) stimuli. Our analyses focused on: (1) ERPs indexing sensory processing and MSI (AV vs. A + V); (2) α-ERD as a measure of cortical excitability; and (3) theta-band functional connectivity using the weighted phase-lag index (wPLI) to quantify coordination between sensory and frontal regions. By integrating these complementary measures, we aimed to delineate the mechanisms underlying reduced multisensory facilitation in autism and to test whether such deficits arise from altered sensory encoding, impaired cortical engagement, disrupted network-level coordination, or a combination thereof. Including unaffected siblings further enabled us to probe the potential genetic underpinnings of these neural phenotypes.

## Methods

### Participants

The study initially included 38 non-autistic (NA), 54 autistic (AU), and 28 sibling (SIB) participants, all between 8 and 13 years of age. After excluding participants based on poor behavioral performance, EEG and eye-tracking data quality, and gaze position for visual stimulation conditions (see corresponding sections for data exclusion criteria), the final analyses were conducted on 36 NA, 40 AU, and 26 SIB participants (see Table 1 for participant characteristics). To be included in the AU group, participants had to meet diagnostic criteria for AU on the basis of the following measures: *1*) autism diagnostic observation schedule 2 (ADOS-2) (Lord et al., 1994); *2*) diagnostic criteria for autistic disorder from the *Diagnostic and Statistical Manual of Mental Disorders* (DSM-5 (American Psychiatric, 2022)); *3*) clinical impression of a licensed clinician with extensive experience in diagnosis of children with ASD. Due to precautions during the COVID-19 pandemic, a subset of AU participants (n=9) was not able to complete the ADOS-2 (Lord et al., 1994), as masking requirements impacted administration. These participants instead underwent the Childhood Autism Rating Scale 2 (CARS-2) and Autism Diagnostic Interview-Revised (ADI-R) (Rutter et al., 2003) for diagnostic assessment. Participants in the NA group met the following inclusion criteria: no history of neurological, developmental, or psychiatric disorders, no first-degree relatives diagnosed with AU, and enrollment in age-appropriate grade in school. SIB group participants met the same criteria as the NA group, except that they had a sibling diagnosed with AU. Exclusion criteria for all groups included: (1) a known genetic syndrome associated with an IDD (including syndromic forms of AU), (2) a history of or current use of medication for seizures in the past 2 years, (3) significant physical limitations (e.g., vision or hearing impairments, as screened over the phone and on the day of testing), (4) premature birth (<35 weeks) or having experienced significant prenatal/perinatal complications, or (5) a Full Scale IQ (FS-IQ) of less than 80. All procedures were approved by the Institutional Review Board of the Albert Einstein College of Medicine and adhered to the ethical standards outlined in the Declaration of Helsinki. All participants assented to the procedures, and their parents or guardian signed an informed consent approved by the Institutional Review Board of the Albert Einstein College of Medicine. Participants received nominal recompense for their participation (at $15 per hour).

**Table 1.** Demographic characteristics. NA=non-autistic control group; AU=autistic group; SIB=unaffected siblings of autistic individuals.

|  | NA (n=36) | AU (n=40) | SIB (n=27) | Statistic | p-value |
| --- | --- | --- | --- | --- | --- |
| Sex<br>F\|M | 17 \| 19 | 7 \| 33 | 16 \| 11 | - | - |
| Age<br>Mean±Std<br>(range) | 10.43 ± 1.8<br>(8-13) | 10.77 ± 1.56<br>(8-13) | 10.6 ± 1.42<br>(8-13) | F=0.397 | 0.674 |
| FSIQ | 103.97 ± 14.52 | 97.10 ± 15.87 | 102.6 ± 15.89 | F=2.03 | 0.137 |
| ADOS | - | 7.63 ± 1.70 | - | - | - |
| F-score | 0.92 ± 0.06 | 0.89 ± 0.08 | 0.90 ± 0.08 | F=1.10 | 0.33 |

### Experimental procedure

Participants were seated in a chair in an electrically shielded room (International Acoustics Company, Bronx, New York), 70 cm away from the visual display (Dell UltraSharp 1704FPT). The stimuli, controlled by Presentation software (Neurobehavioral Systems), included three types: a red disc (’Visual’), a 1000Hz tone (’Audio’), and their simultaneous presentation (’Audiovisual’). Participants were instructed to press a button as quickly as possible upon detecting any stimulus. The auditory stimulus was 1000Hz, 60ms tone presented binaurally (75 dB SPL). The visual stimulus was a red disc subtending to 1.5 degrees, displayed above a fixation cross. The audiovisual stimulus was a simultaneous presentation of both. Each trial presented a pseudo randomly chosen stimulus (A, V, or AV; represented equiprobably), with stimuli delivered through headphones (HD 650 Sennheiser) and displayed on a flat-panel LCD (Dell UltraSharp 1704FPT, 60Hz). A jittered randomly sampled interstimulus interval (1000-3000ms) reduced onset predictability and therefore anticipatory responses in the baseline period. The task consisted of 400 trials across 4 blocks (100 trials per block), each block lasting approximately 3 minutes and 40 seconds. Button presses were recorded using a response pad (Logitech Wingman Precision Gamepad). Triggers indicating stimulus onset were sent to the EEG stimulus channel from the PC acquisition computer via Presentation software.

### Behavioral data

The following performance metrics were calculated for each participant: Miss rate: proportion of missed responses (*false negatives / total stimuli*). Precision: accuracy of correct responses (*true positives / (true positives + false positives*)). Hit rate: proportion of correct detections (*true positives / (true positives + false negatives*)). F-score: combined measure of precision and Hit rate (*2 × (precision × Hit rate) / (precision + Hit rate*)). Three participants from the ASD group were excluded due to non-compliance with the task, defined as having a miss rate exceeding 50% in at least one of the three stimulation conditions.

### Race model analysis

To determine whether multisensory benefits exceeded statistical facilitation predicted by Raab’s independent race model (Raab, 1962) following the approach described in (Crosse et al., 2022), we analyzed cumulative distribution functions (CDFs) of reaction times (RTs) using a stepwise procedure. The analysis was performed on repeat trials (V→V, A→A, AV→AV) since switch trials (A→V, V→A, V→AV, A→AV) introduce additional reaction time costs due to intersensory switching effects (Shaw et al., 2020; Vanneau, Quiquempoix, et al., 2025), which can confound measures of multisensory facilitation. Intersensory switching cost refers to the increased response time observed when the modality of a stimulus changes from one trial to the next (e.g., from auditory to visual), reflecting the cognitive load associated with reorienting attention across sensory modalities (Vanneau, Quiquempoix, et al., 2025). Unisensory trials that were preceded by multisensory trials (AV→A, AV→V) were excluded from the analysis, as they did not cleanly fit within the definitions of repeat or switch trials. The race model was computed separately for repeat and switch trials while incorporating the joint probability term. Specifically, for each participant, we estimated the race model as:

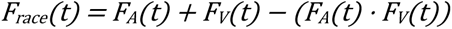

where *F_A_(t)* and *F_V_(t)* represent the CDFs of RTs for the auditory and visual conditions, respectively. The predicted multisensory benefit was calculated as the difference between the race model and the maximum of the two unisensory CDFs at each quantile:

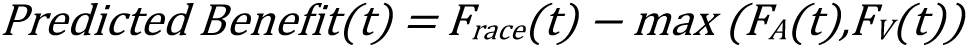

This step ensures that the predicted benefit captures the expected facilitation under a statistical race model assumption. The empirical multisensory benefit was computed by comparing the observed audiovisual (AV) CDF to the maximal unisensory CDF:

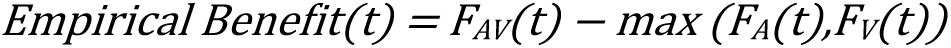

This represents the actual gain in RTs observed during multisensory trials. To test whether the observed multisensory benefit exceeded the statistical prediction, we computed the race model violation as the difference between the empirical and predicted benefits:

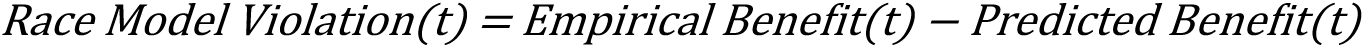

Positive values indicate that the observed multisensory gain is greater than what the race model predicts, suggesting supramodal multisensory integration rather than simple statistical facilitation. As a summary measure of multisensory integration, we quantified the multisensory gain as the total area under the curve (AUC) of the race model violation function:

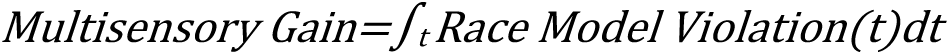

This metric captures the extent to which multisensory RT facilitation exceeds statistical predictions across the RT distribution. By applying this approach separately to repeat and switch trials, we were able to assess whether multisensory integration mechanisms differed based on prior sensory context.

### EEG recordings & preprocessing

EEG data were recorded at a sampling rate of 512Hz using 64 channels BioSemi Active II system (using the CMS/DRL referencing system) with an anti-aliasing filter (−3 dB at 3.6 kHz). Analyses were conducted in python (3.11) using MNE (originally for Minimum Norm Estimation)(Gramfort et al., 2013) and custom scripts available at https://github.com/tvanneau/SFARI-AVSRT. Bad channel detection was performed using the function NoisyChannels (with RANSAC) from the pyprep toolbox (Bigdely-Shamlo et al., 2015). If more than 15% of the channels were detected as bad, the participant was rejected (1 NA, 2 AU and 1 SIB). Bad channels were interpolated using spline interpolation (Perrin et al., 1989). EEG was filtered using a FIR band-pass filter (0.01-40Hz), and Independent Component Analysis (ICA) on 1Hz high-pass EEG was used to identify and manually reject eye-related components (blinks/saccades). Epochs were created from -500 to +800ms around each stimulation with a baseline-correction from -50ms to +20ms stimulus onset and referenced to a common average reference. EEG trials were only included if the participant responded within 100 to 1500 ms after stimulus onset. Trials where a participant provided more than two responses were also removed to prevent random over-clicking behaviors. To further ensure EEG data quality, eye-position filtering was applied as described below.

### Eye-tracking

Gaze position and pupil data were recorded with an EyeLink 1000 (SR Research Ltd., Mississauga, Ontario, Canada) at a sampling rate of 500 Hz. Due to the absence of eye-tracking recordings, caused by hardware issues or poor signal quality due to glasses, nine participants in the AU group and one in the SIB group were excluded from all analyses. Gaze position data were used to filter out ‘Visual’ and ‘Audiovisual’ EEG trials where the participant was not fixating within a defined region of interest (ROI) centered on the fixation cross (±7° from fixation in the x-axis and ±5° on the y-axis). This filtering was applied over a 40 ms window around stimulus onset (-20 ms to +20 ms). The proportion of EEG trials rejected due to eye-tracking filtering was as follows: NA group: 14.23% ± 2.12 (Visual), 15.08% ± 2.13 (Audiovisual). AU group: 20.57% ± 2.85 (Visual), 20.59% ± 2.77 (Audiovisual). SIB group: 17.14% ± 3.70 (Visual), 19.38% ± 3.73 (Audiovisual).

After behavioral, eye-tracking and artifact filtration, the final number of EEG trials for each participant were, on average: 96.61 ± 4.8 for visual, 111.56 ± 3.15 for auditory, 100.53 ± 4.02 for audiovisual for the NA group; 84.92 ± 4.73 for visual, 101.62 ± 4.00 for auditory, 89.28 ± 4.06 for audiovisual for the AU group; 89.12 ± 5.68 for visual, 108.62 ± 4.60 for auditory, 95.19 ± 5.36 for audiovisual for the SIB group.

### ERP analysis-Unisensory processing

Group differences in auditory and visual sensory processing were first assessed. To quantify ERP amplitude and latency in response to visual stimulation, we selected a cluster of six electrodes (PO3, PO7, O1, O2, PO4, PO8) over parieto-occipital scalp where the VEP is greatest (Luck et al., 2000; Molholm et al., 2002). Amplitudes were calculated as the mean signal within defined temporal windows, while latencies were extracted by identifying the local maximum or minimum peak (depending on the component polarity) within the same window. The temporal windows were defined based on the literature and adjusted, if necessary, based on the timing of the components in the grand average, without regard to group. The P1 and N1 components of the visual ERP were analyzed in the time ranges of 120–170 ms and 190–230 ms, respectively (Curran et al., 2001; Di Russo et al., 2002; Vanneau, Quiquempoix, et al., 2025). For the auditory ERP, we analyzed responses from a cluster of two right and two left temporo-parietal electrodes (T7, TP7, T8, TP8), where auditory responses were expected to be strongest for this age group (Gomes et al., 2001; Ponton et al., 1996), and as confirmed by inspection of the data. The auditory N1 and P2 were extracted within the time windows of 120–170 ms and 220–350 ms, respectively (Fitzroy et al., 2015; Liasis et al., 2003).

### ERP analysis-Multisensory processing

To quantify multisensory integration (MSI) in the ERPs, we compared the audiovisual (AV) response to the linear sum of the unisensory auditory (A) and visual (V) responses (Molholm et al., 2002). Any divergence from the linear sum during the sensory processing timeframe (<200 ms post stimulus onset) is interpreted as evidence that the inputs were processed differently when they came together, i.e., that MSI occurred. For each participant and electrode, we computed a difference wave:

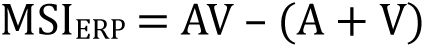

The generators and latencies of MSI effects are not readily constrained to a small set of scalp regions or time windows (Senkowski & Engel, 2024). Therefore, cluster-based permutation testing in which sensor adjacency and temporal contiguity are leveraged to identify significant clusters while controlling for multiple comparisons (Maris & Oostenveld, 2007) was used as an unbiased approach to test for evidence of MSI across all electrodes and time-points within the time-window of consideration. This analysis was constrained to the first 250 ms post-stimulus onset since later cognitive components (such as the P3) would be represented twice in the summed response, leading to an artificial MSI effect.

### Spectral analysis

Spectral analysis was performed using complex Morlet wavelet convolution (tfr_morlet function in MNE) between 2 and 40Hz with a FWHM in the spectral domain of 3.25Hz and 187ms in the temporal domain (Cohen, 2014) that correspond to: number of cycles = frequency / 2 (with frequency as a vector of frequency from 2 to 40 in steps of 0.19Hz). The non-phase locked power was calculated after subtracting the ERP from each trial for each individual (Kalcher & Pfurtscheller, 1995). Power values for clusters of channels where the auditory and visual sensory evoked ERPs were measured were averaged (i.e. a bilateral parieto-occipital cluster for the visual response and bilateral temporal clusters for the auditory evoked response). For the total power and the non-phase locked power two different baseline normalization methods were tested since baselining method can influence the results (Grandchamp & Delorme, 2011): A trial-by-trial baseline normalization and an average baseline normalization was applied after averaging all the trials. Both methods were performed independently for every individual using a -200ms to 0ms baseline period and a decibels (dB) normalization (10log10(Post-stim power/Baseline power). The baseline approach did not influence the data; the results and figures are from data using the average baseline normalization approach.

### Alpha – Multisensory integration analysis

To test MSI effects in the alpha band, we compared α-ERD to audiovisual (AV) stimuli with the linear sum of the unisensory responses (A+V). Because linear summation is not valid in the spectral domain (Senkowski et al., 2006), A and V epochs were first summed in the time domain on a trial-by-trial basis; alpha-band power was then extracted from these A+V waveforms using the same Morlet-wavelet parameters described above. We generated a bootstrap distribution of A+V spectral values (1000 resampling trials with replacement) and z-scored the AV alpha power against this distribution for each group. α-ERD was quantified by averaging alpha power over time (200-500ms) within a parieto-occipital electrode cluster defined from the grand-average topography (collapsed across groups/conditions) and grounded in prior work showing maximal alpha suppression over visual cortex during audiovisual processing.

### Connectivity Analysis

Functional connectivity was assessed using the weighted Phase Lag Index (wPLI), computed with the spectral_connectivity_epochs function from MNE. To improve spatial specificity and reduce volume conduction effects, data were first spatially filtered using a Laplacian transform. Focusing on Theta and Alpha bands, wPLI was calculated in the 3–13 Hz frequency range using complex Morlet wavelets, with the number of cycles increasing linearly from 2 cycles at 2 Hz to 10 cycles at 13 Hz (incremented by 0.1 cycle per frequency step).For each stimulation condition, wPLI values were computed for all pairs of EEG channels, and connectivity between parietal and frontal regions was quantified by averaging wPLI values across all corresponding electrode pairs (Figure 5). We calculated percentage change of wPLI from a pre-stimulus baseline window (-200 to - 50 ms) for the averaged parietal–frontal connectivity within each group. It is important to note that traditional MSI testing (Av versus A+V) as done for the ERPs and for alpha is not possible here because wPLI is a phase-based measure of *inter-regional coupling* derived from the cross-spectrum of two signals. Linearly summing A and V within each channel to form (A+V) would corrupt the phase-relationship information across sites, producing terms that have no interpretable relation to “unisensory coupling.” Therefore, comparison of connectivity by stimulus condition and group will be used to understand whether there are multisensory related differences in functional connectivity in ASD.

### Statistical analysis

All statistical analyses were conducted using Jamovi (The jamovi project, version 2.3.28, https://www.jamovi.org) for linear mixed models and Python (MNE-Python) for permutation-based testing. Comparisons of RTs across groups and sensory modalities were performed using Linear Mixed Models (LMMs) to account for individual variability. Age was included as a covariate, and all potential interactions were tested. We report the F-score from the fixed-effects omnibus test, along with partial Eta Squared (η^²^p) as an effect size measure and the corresponding p-value (α = 0.05). The Shapiro-Wilk test was used to assess the normality of residuals. Statistical significance for the race model analysis was evaluated using nonparametric permutation tests (Maris & Oostenveld, 2007), implemented in Python using the permutation_t_test function from MNE. A one-sample t-test (1,000 permutations) was performed to determine whether the observed race model violation (empirical minus predicted benefit) was significantly greater than zero within groups. To control family-wise error rate, a t_max_ correction was applied (Blair et al., 1994; Thomas et al., 1994). For race model violation, we used two subject-level metrics: (i) the maximum violation within the first four RT quartiles (per subject), and (ii) the area under the entire violation curve (Crosse et al., 2022). Each metric was then compared across groups using one-way ANOVA. For within-condition ERP amplitudes, we tested normality with Shapiro–Wilk and, when assumptions were met, ran one-way ANOVA with group as a fixed factor. For ERP multisensory integration (MSI), we employed two complementary approaches: an unbiased spatio-temporal cluster-based permutation test (Maris & Oostenveld, 2007) evaluating all sensors and time points while controlling the family-wise error rate; and a ROI/time-window analysis extracting the AV − (A+V) difference between 145–165 ms for each participant, followed by one-way ANOVA across groups. For induced power and connectivity (wPLI), we used the same two-tier strategy: nonparametric temporal cluster-based permutation tests (1,000 permutations; α = 0.05) without a priori temporal selection; and a classical peak amplitude/latency detection per subject (or averaging over a time-window for wPLI) with subsequent one-way ANOVA for group differences. Because alpha ERD exhibits a clear trough with interpretable timing, we summarize it with peak amplitude and latency; in contrast, time-resolved wPLI is noisier and lacks a sharp, physiologically interpretable peak, so we use a priori window means (supported by cluster-based tests) to obtain reliable group estimates. The specific MSI procedure for alpha ERD is detailed in “Alpha—multisensory integration analysis. Finally, baseline relative alpha power was analyzed with one-way ANOVA (α = 0.05) after confirming normality of the distributions.

## Results

### Behavior: Reduced multisensory gain in ASD and SIB groups

The results demonstrate a main effect of sensory modality on reaction time (Figure 1B; *F* = 145.7, η^²^p = 0.59, *p* < 0.001), with audiovisual stimulation leading to faster reaction times compared to both auditory (ß 83.21 ms, *t* = 16.57, *p* < 0.001) and visual conditions (ß = 53.82 ms, *t* = 10.72, *p* < 0.001). Additionally, reaction times were significantly faster for the visual modality compared to the auditory modality (ß = 29.4 ms, *t* = 5.86, *p* < 0.001). A significant effect of age was observed (*F* = 33.83, η^²^p = 0.25, ß = 31.66 ms, *p* < 0.001), along with a significant interaction between age and sensory modality (*F* = 4.1, η^²^p = 0.04, *p* = 0.018) due to a reduced benefit of increasing age for the visual modality (Supplementary Figure 1). No significant main effect of group or interactions between group, age, or sensory modality were found (Main group effect: *F* = 0.4, η^²^p < 0.01, *p* = 0.66; Group × Sensory Modality: *F* = 0.19, η^²^p < 0.01, *p* = 0.94; Group × Age: *F* = 1.90, η^²^p = 0.03, *p* = 0.154).

**Figure 1.**
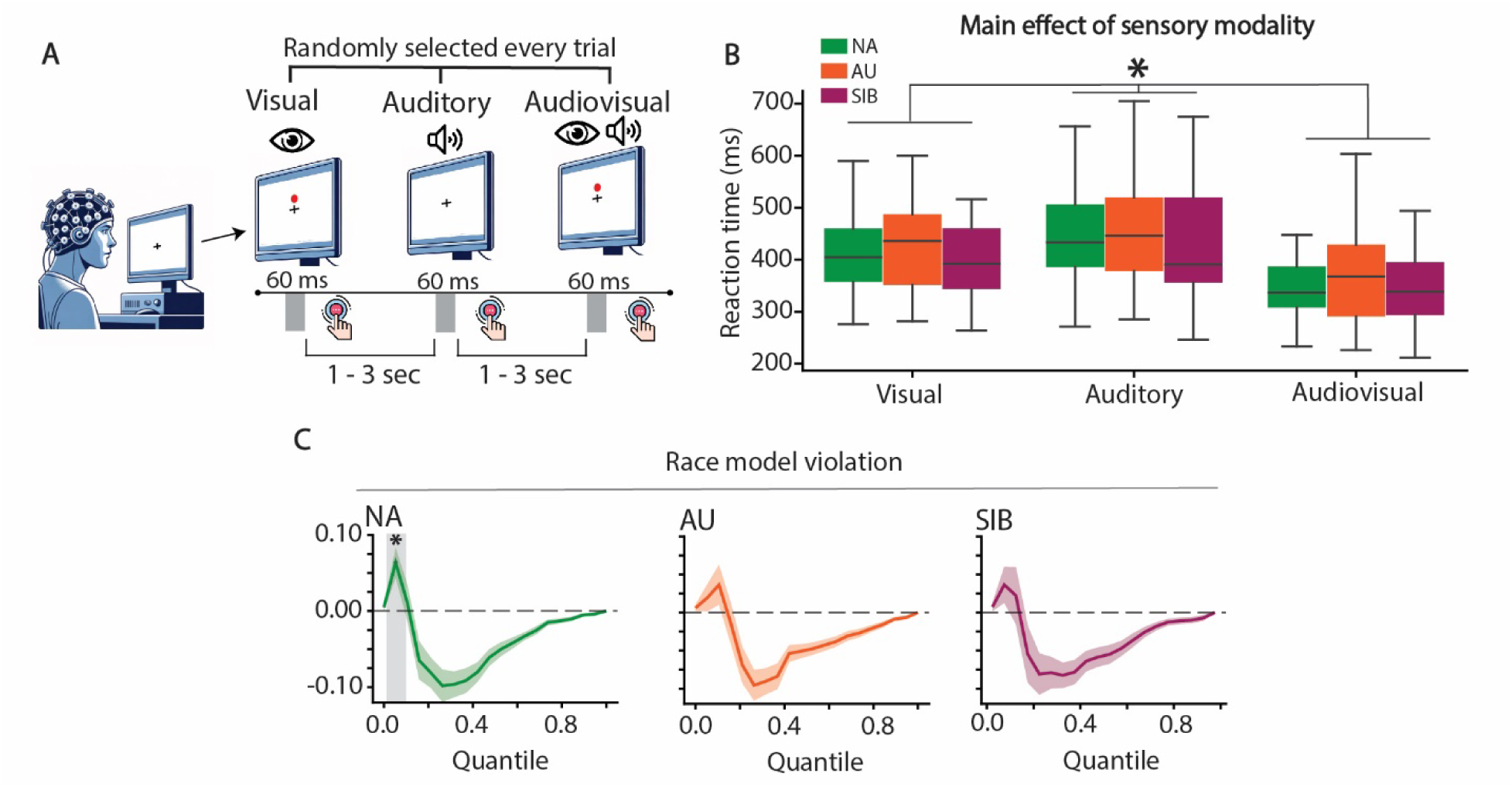
Comparable RTs between groups but reduced multisensory gain for AU and SIB groups. **(A)** Experimental paradigm: Participants were instructed to press a button as quickly as possible upon detecting any stimulus type. **(B)** Reaction times (RTs) across sensory modalities and groups: Mean RTs (in ms) for Non-autistic (NA, green), Autistic (AU, orange), and Siblings of autistic children (SIB, purple). Asterisks (*) indicate statistically significant differences determined by a linear mixed model (α = 0.05). **(C)** Race model violation curves: Difference between empirical and predicted benefits (see Methods) in NA (green), AU (orange), and SIB (purple). Shaded areas indicate the standard error of the mean (SEM). Asterisks (*) denote significant differences identified via permutation testing using a one-sample t-test with tmax correction.

Within-group, race model violation was detected in the NA group, whereas no violation emerged for AU or SIB (Figure 1C; t-test-based permutation statistics, *t*-max correction). Between-group comparisons were not significant for either summary metric, the maximum violation within the first four RT quartiles or the AUC of the full violation curve (Supplementary figure 1; F_s_ < 0.15 and p_s_ > 0.8).

### ERPs: Similar unisensory evoked responses between groups

The three participant groups exhibited similar topographical distributions and characteristic ERP waveforms for each sensory modality (Figure 2A). The visual response was focused over occipital scalp and comprised of a positive-going response peaking at 145 ms followed by a negative-going response peaking at 215 ms, corresponding to the P1 and N1 components respectively. The auditory response was focused over fronto-central and temporal scalp regions, reflecting the dipolar configuration generated by auditory cortices, with classic P1 and N1 responses that peaked at 80 and 120 ms respectively, followed by the P2 peaking at about 180 ms. Waveforms are displayed for temporal channels where the so-called N1 complex response was greatest (as expected in this age range) and from where measurements for statistical tests were taken. Peak amplitude and corresponding latency values for the sensory components were extracted for each component of interest (see Methods). The amplitude of the visual P1 did not differ significantly between groups, although there was a trend toward a higher amplitude in the NA group (Figure 2B; *F* = 2.93, η^²^p = 0.05, *p* = 0.058). Similarly, P1 latency was not significantly different between groups (*F* = 0.62, η^²^p = 0.01, *p* = 0.53). No group differences were found for the amplitude or latency of the visual N1 (Amplitude: *F* < 0.01, η^²^p < 0.01, *p* = 0.9; Latency: *F* < 0.01, η^²^p < 0.01, *p* = 0.9). For the auditory ERP, neither the N1 or P2 components significantly differed by group in either amplitude (*N1*: *F* = 0.15, η^²^p < 0.01, *p* = 0.86; *P2*: *F* = 1.77, η^²^p = 0.03, *p* = 0.17) or latency (*N1*: *F* = 1.06, η^²^p = 0.02, *p* = 0.34; *P2*: *F* = 0.56, η^²^p = 0.01, *p* = 0.56). Notably, when we compared the ERPs without the eye-tracking filtration (see Methods), statistical differences emerge for the visual P1 with a main group effect (Supplementary Figure 2; F = 3.12, η^²^p = 0.06, p = 0.04). This emphasizes the importance of carefully controlling gaze when comparing responses to visual stimuli across groups.

**Figure 2.**
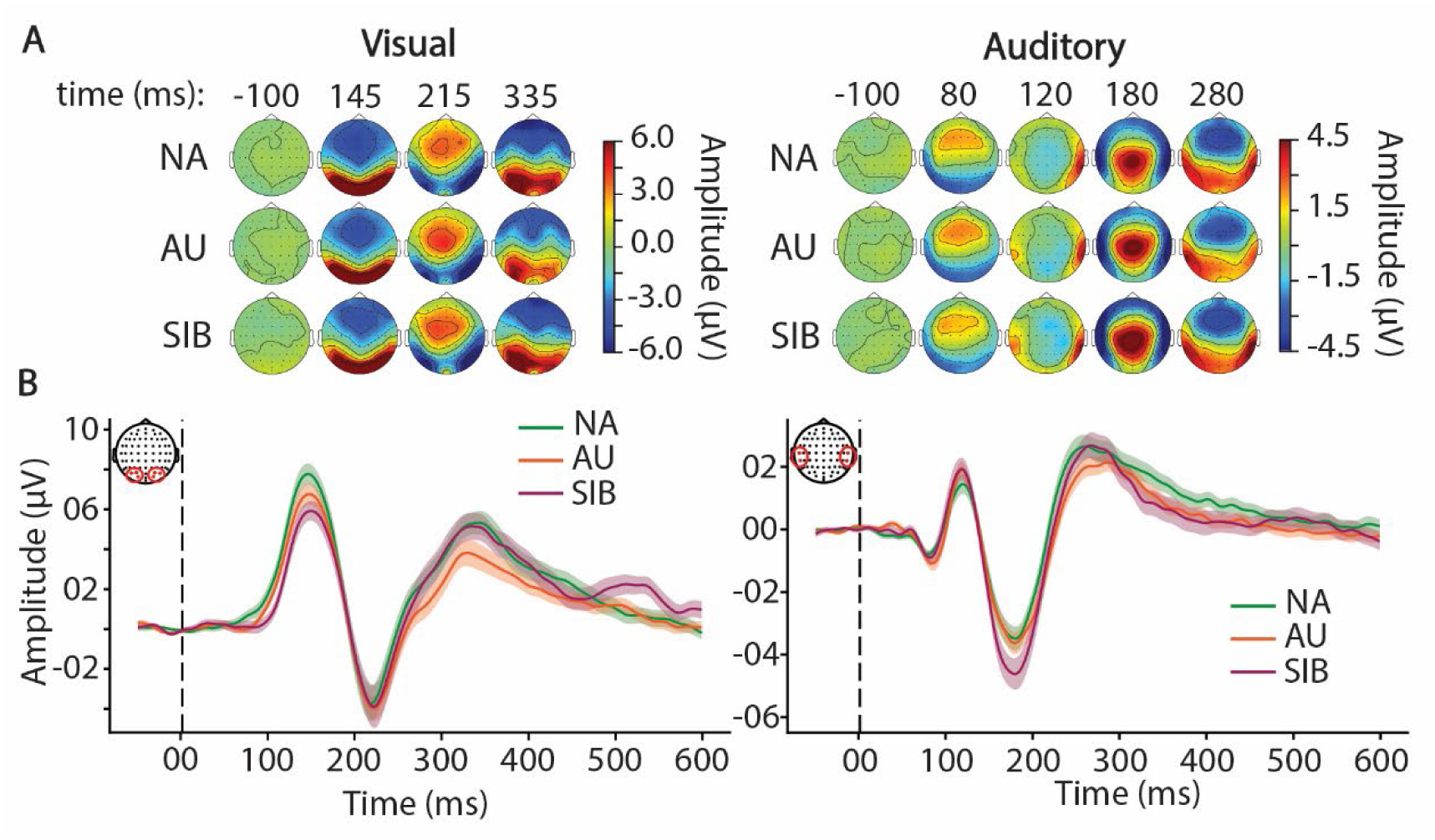
Similar unisensory ERPs among groups. (**A**) Topographical distribution of event-related potentials (ERPs) in response to visual stimulation (left) and auditory stimulation (right), displayed at the timing of key ERP components for each group (non-autistic [NA]: top row; autistic [AU]: middle row; siblings of autistic individuals [SIB]: bottom row). (**B**) Averaged visual (left) and auditory (right) ERPs for a cluster of occipital channels for visual, and a cluster of temporal channels for auditory, in NA (green), AU (orange) and SIB (purple). The shaded areas indicate the standard error of the mean (SEM) in the corresponding color.

### ERPs: MSI is reduced in autistic and sibling groups

We next contrasted the audiovisual (AV) ERP with the linear sum of the unisensory auditory and visual ERPs (A+V). Unlike the preceding analyses of the unisensory responses that used predefined latencies and electrode clusters based on well-known component characteristics, the underlying generators and resulting dipolar configurations of MSI effects are not well characterized and could plausibly arise at multiple latencies and scalp sites. Therefore, using an unbiased approach, a spatio-temporal cluster-based permutation test (Maris & Oostenveld, 2007) was applied to evaluate MSI at all sensors and time points while controlling the family-wise error rate. Tests were restricted to the <250 ms time window to minimize contributions from later, non-sensory components (e.g., P3), whose nonlinearity could lead to an artificially inflated summed (A+V) response. In the NA group, this analysis revealed a significant parieto-central cluster from ∼110–180 ms (Figure 3A); no significant clusters were found in the AU or SIB groups. For visualization, we then averaged ERPs across the electrodes comprising the NA cluster, which suggested a latency difference between AV and A+V specific to the NA group (Figure 3B). A post-hoc test in which the average value of the difference wave (AV – (A+V)) between 145 to 165 ms was compared across groups (for the cluster of significant channels identified with the spatio-temporal approach, illustrated in Figure 3B) revealed a significant effect of group (F = 6.28, η^²^p = 0.14, p = 0.003), with higher values for NA compared to both AU (t = 2.9, p = 0.013) and SIB (t = 3.16, p = 0.006) without differences between AU and SIB (t = 0.58, p = 0.83).

**Figure 3.**
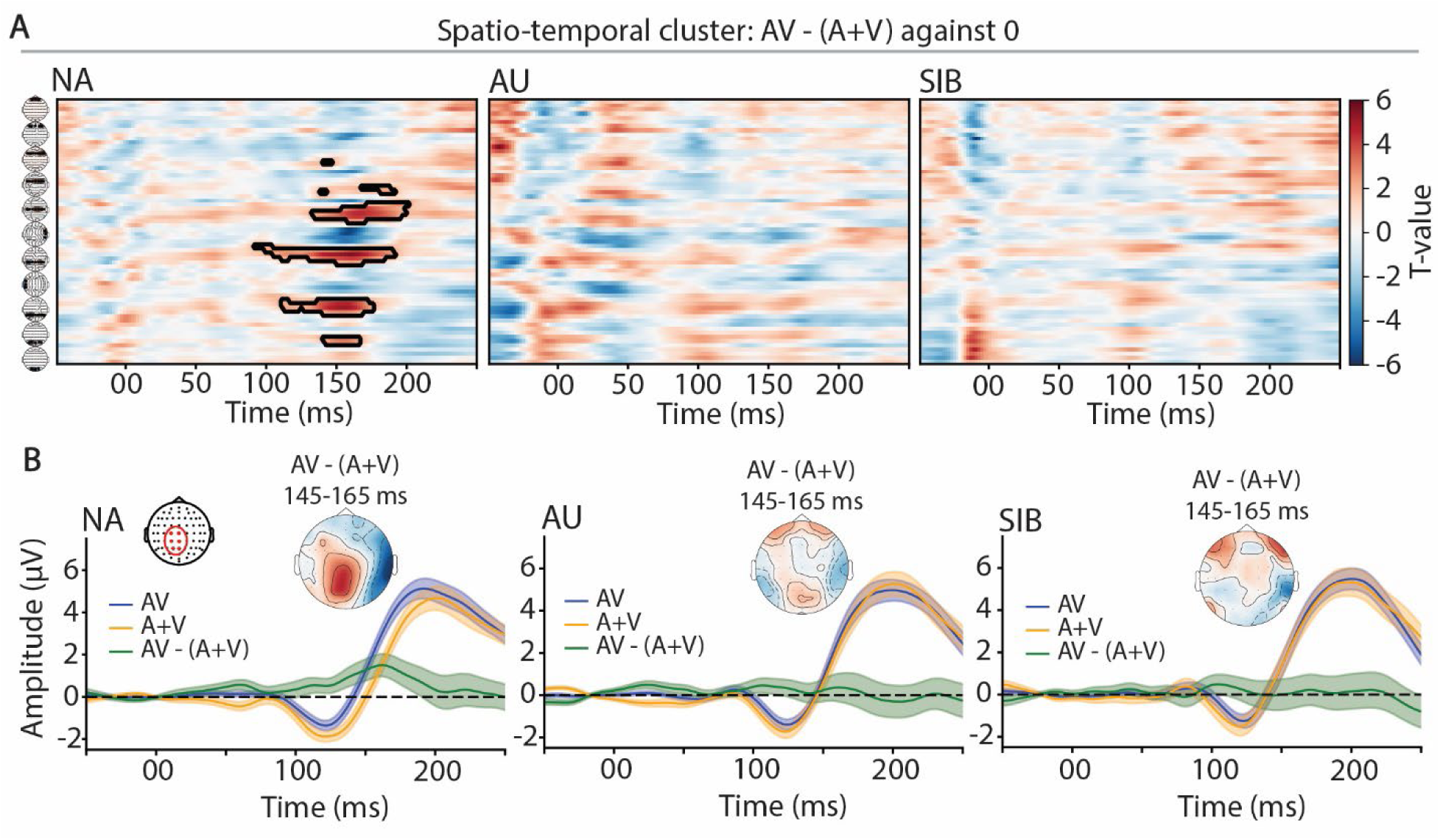
Lack of MSI effects in autistic and sibling groups. (**A**) Spatio-temporal cluster plot of T-values (one-sample test against 0) for the difference (AV – [A+V]), with significant clusters outlined in black. Plots are shown separately for NA (left), AU (middle), and SIB (right) groups. (**B**) Grand-average ERPs over a cluster of left parieto-central electrodes (selected from the significant spatial cluster of the permutation statistics for the NA group) in response to audiovisual stimuli (blue), the sum of unisensory stimuli (orange), and their difference (green). The inset shows the topographic distribution of the difference wave (AV – [A+V]) averaged between 145–165 ms, displayed for the NA group (left), AU group (middle), and SIB group (right).

### Neuro-oscillations: Similar α-ERD in response to unisensory in all groups

We examined spectral activity in the alpha (7-13 Hz) frequency band. Analysis of induced alpha activity demonstrated a pronounced decrease in power (e.g., α-ERD) centered over occipital channels in response to visual stimulation (Figure 4A) and a subtle decrease over lateral temporal regions in response to auditory stimulation. Auditory-driven α-ERD was notably weaker than visual-driven ERD (Figure 4B). Temporal cluster-based permutations between each group in each sensory condition did not reveal significant differences between any groups for both sensory conditions, as with the ANOVA performed on peak amplitude values and the associated latency (Supplementary Figure 3; p_s_ > 0.15). Because the spectral power was baseline-normalized, another analysis was performed to assess whether the absence of group differences in α-ERD could be attributed to differences in pre-stimulus alpha power (i.e., lower pre-stimulus alpha could lead to lower alpha desynchronization since it is a ratio from the baseline). We measured relative alpha power during the baseline period and found no support for significant differences between groups (Figure 4C; Visual: *F* = 0.27, η^²^p < 0.01, *p* = 0.76; Auditory: *F* = 0.64, η^²^p = 0.01, *p* = 0.52).

**Figure 4.**
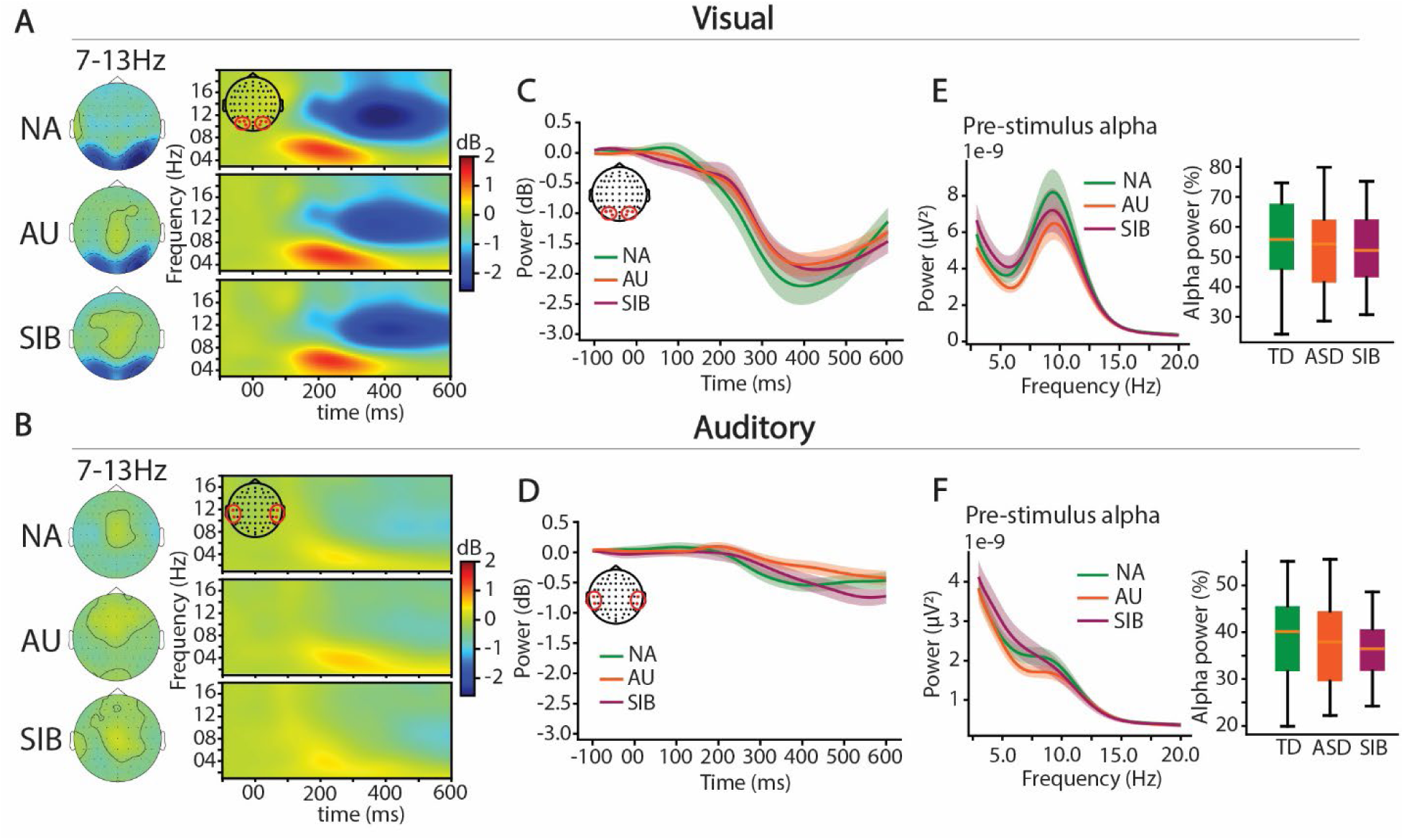
Similar α-ERD to unisensory stimulation. (**A**) Averaged topographical distribution of induced alpha activity (7–13 Hz) in response to visual stimulation along with the corresponding time-frequency map for a cluster of occipital channels in response to the visual stimulation for NA (top row), AU (middle row), and SIB (bottom row). (**B**) same as A for auditory stimulation and a cluster of temporal electrodes (see topographical illustration). (**C**) Induced alpha activity averaged over a cluster of channels (see corresponding topomaps) for NA (green), AU (orange), and SIB (purple) in response to the visual stimulation and (**D**) in response to the auditory stimulation. (**E**) Power spectrum density and relative alpha power for the pre-stimulus baseline period (-200ms to stimulus onset) in response to the visual and (**F**) to the auditory stimulation for NA (green), AU (orange) and SIB (purple). The shaded areas in the plots represent the standard error of the mean (SEM) in the corresponding colors.

**Figure 5.**
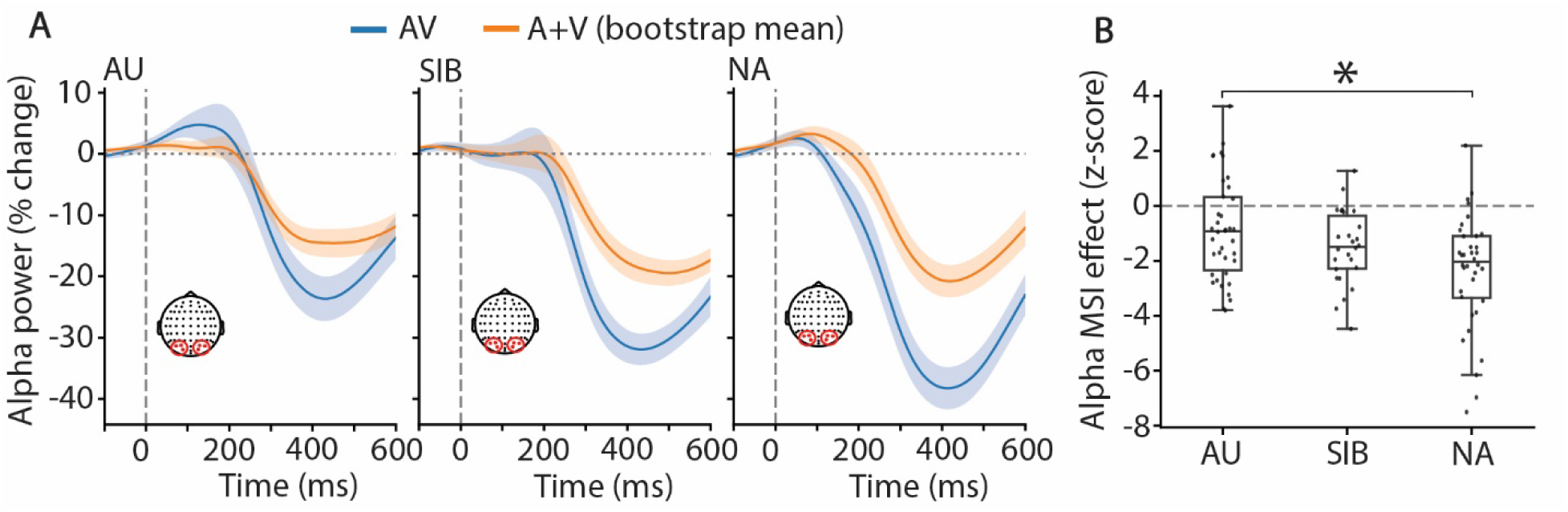
Attenuated multisensory enhancement of α-ERD in autism. (**A**) Induced alpha activity averaged over a cluster of parieto-occipital channels (see corresponding topomaps) in response to audiovisual stimulation (blue) and for A+V prediction (orange). The A+V prediction was generated by pairing and summing A and V trials in the time domain (bootstrapped pairings) before time–frequency decomposition. Traces show group means (AU left, SIB middle, NA right) from −100 to 600 ms. (**B**) MSI z-scores computed per participant as the ROI alpha power (200–500 ms) in AV minus the mean of the bootstrapped A+V distribution, divided by its SD. Asterisks indicate significant post-hoc group differences following a main ANOVA effect (α = 0.05). All groups showed a significant MSI effect relative to zero (one-sample t-test).

### Neuro-oscillations: α-ERD MSI reduced in autistic and sibling groups

To determine if AV α-ERD differed from the sum of the unisensory α-ERDs, we compared α-ERD in response to audiovisual (AV) stimuli with the linear summation of the unisensory auditory and visual conditions (A+V). A bootstrapping procedure was applied to the time-domain A+V trials to generate a distribution of spectral values, and the AV alpha power was subsequently z-scored against this distribution for each group (see Methods). α-ERD was quantified by averaging power across time (from 200 to 500ms) within the parieto-occipital cluster indicated in Figure 5A. This revealed z-scores significantly different from zero in all groups, indicating a multisensory supra-additive effect (e.g., greater α-ERD; Figure 5B). A one-way ANOVA revealed a significant main effect of group (F = 6.17, η^²^p = 0.113, p = 0.003), with post-hoc tests showing a significant difference between NA and AU (Δ = 1.49, t = 3.51, p = 0.002), but no significant differences between NA and SIB (p = 0.163) or AU and SIB (p = 0.374).

### Reduced Theta-Band Connectivity Between Parieto-Occipital and DLPFC Regions in Autism

To examine whether autistic children show altered long-range functional connectivity between sensory and top-down control regions, we analyzed phase-based connectivity using the weighted Phase Lag Index (wPLI), a measure of the delay in phase between two regions, robust to volume conduction that excludes zero-lag synchronization (Stam et al., 2007). By focusing on data from channels over frontal and occipital scalp, this analysis addressed connectivity between the parieto-occipital cortex and the dorsolateral prefrontal cortex (DLPFC), a region well established in supporting top-down attentional control (Katsuki & Constantinidis, 2012; Knight et al., 1995). Time-frequency analysis revealed a similar increase in wPLI across groups in response to visual stimulation, with connectivity predominantly in the upper theta band (6–7 Hz) and peaking around 220 ms post-stimulus (Figure 6A-B). Auditory stimulation elicited only minimal modulation of frontal to parieto-occipital connectivity, with no differences observed between groups. By contrast, during audiovisual processing AU children showed significantly reduced connectivity between parieto-occipital and frontal regions from ∼110–280 ms post-stimulus, relative to NA peers (Figure 6B). No significant differences emerged for NA vs. SIB or AU vs. SIB. To complement this permutation approach we also extracted the average wPLI value (150-300ms) for each subject in each group and performed an ANOVA, which revealed a significantly higher wPLI for the NA group compared to the AU group in the audiovisual condition (F(2,99) = 3.17, η^²^p = 0.06, p = 0.04; NA vs AU: t = -2.5, p = 0.03; NA vs SIB: t = -0.9, p = 0.61; AU vs SIB: t = -1.3, p = 0.38). Importantly, no group differences were observed in induced theta power over fronto-central regions (Supplementary figure 4), suggesting that the wPLI differences cannot be explained by local variations in signal strength. While wPLI is primarily a phase-based index, its weighted formulation includes amplitude information, making this control analysis essential. Applying the same analysis to data from channels over lateral temporal and parieto-occipital scalp regions (where the auditory and visual responses are greatest) during unisensory and audiovisual stimulation revealed comparable connectivity profiles across groups, consistent with intact coupling between sensory regions in ASD (Supplementary Figure 5).

**Figure 6.**
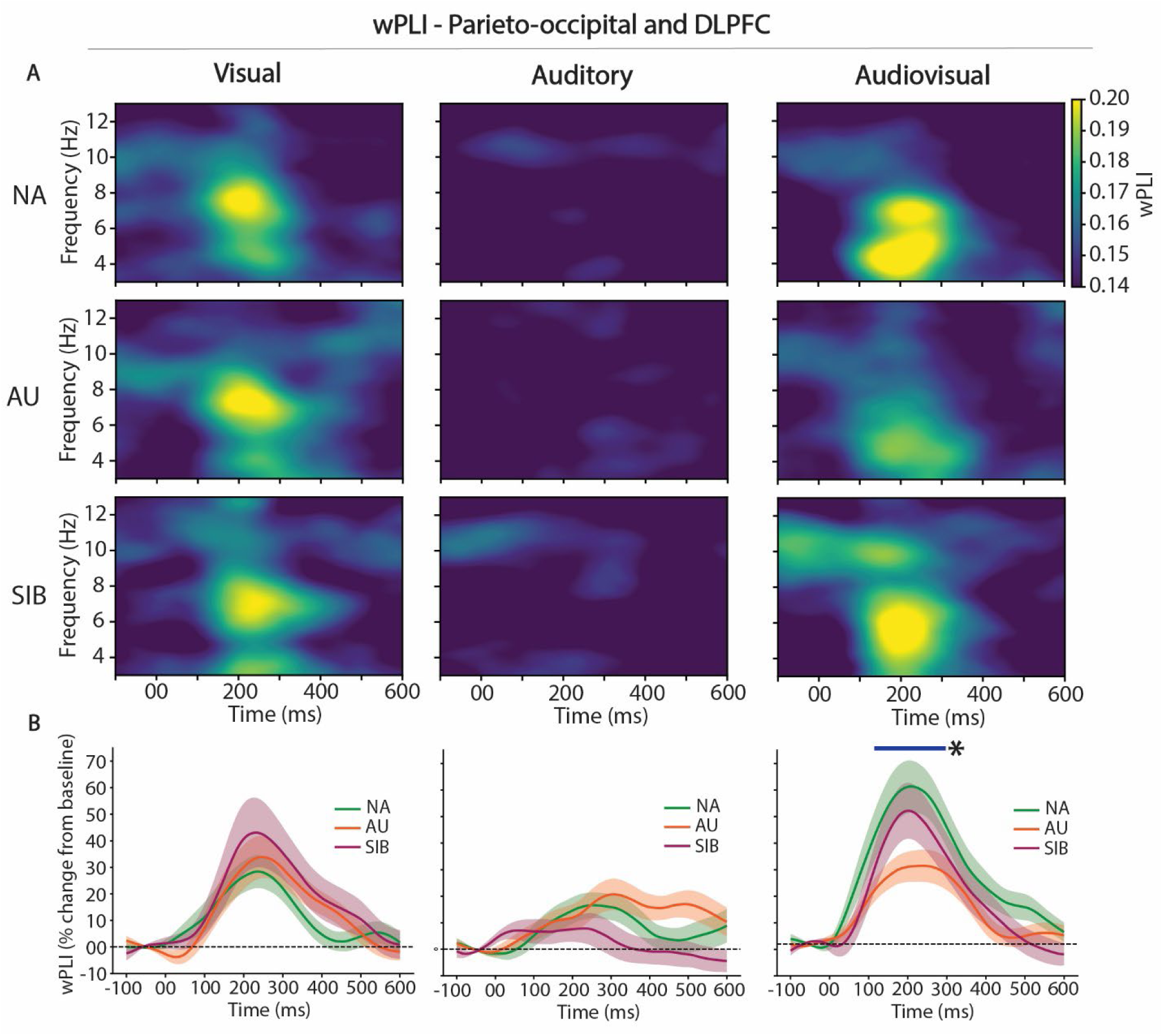
Evidence for Reduced Theta-based connectivity between parieto-occipital and DLPFC in response to audiovisual stimulation in autism. (**A**) Time-frequency representation of the weighted-phase lag index (wPLI) between a cluster of parieto-occipital channels and a cluster of channels located at the position of the dorsolateral prefrontal cortex (DLPFC) in response to a visual stimulation (left), auditory stimulation (center) and audiovisual (right) for the NA group (top row), AU group (middle line) and SIB group (bottom line). (**B**) wPLI averaged for the theta band (4-7Hz) for NA (green), AU (orange), and SIB (purple) for the corresponding sensory stimuli. The blue bar (*) indicates statistically significant differences (between NA and AU group) determined by cluster-based permutation statistics (α = 0.05).

## Discussion

### Converging evidence for reduced multisensory related cortical activation and fronto–parietal coordination in autism

Impaired multisensory processing in children with autism is a robust finding, yet the underlying neural mechanisms are not well understood. Here, neurophysiological and behavioral data recorded from children with and without autism while they engaged in a classic multisensory task (Brandwein et al., 2013; Molholm et al., 2002) were analyzed to test brain mechanisms that might underlie altered MSI in autism. Replicating prior findings (Brandwein et al., 2013; Crosse et al., 2022), the behavioral data revealed reduced MSI in autistic children. Furthermore, this multisensory facilitation of behavior was also reduced in unaffected siblings (SIB) of autistic individuals, showing for the first time that altered MSI in autism may reflect inherited neurobiological differences.

Guided by evidence that early MSI arises in or near sensory cortices (Brandwein et al., 2013; Mercier et al., 2013; Molholm et al., 2002) and by models implicating higher-order control regions (Cuppini et al., 2017; Molholm et al., 2006; Monti et al., 2023; Todd & Bahrick, 2023), we focused analyses on sensory cortical responses and fronto–posterior connectivity. Three main neural findings emerged. First, multisensory alteration of the ERP was only detected in the non-autistic (NA) group, over parieto-occipital scalp. Second, multisensory enhancement of alpha-band event-related desynchronization (α-ERD) was attenuated for autistic (AU) participants (but not SIBs) compared to NA. Third, theta-band wPLI revealed reduced coupling between frontal and posterior regions during AV processing in AU, whereas connectivity for unisensory trials did not differ across groups.

In NA children, the parietal MSI effect aligns with prior reports of superadditive AV effects over left parietal sites ∼100–150 ms (AV > A+V; (Brandwein et al., 2013; McCracken et al., 2019; Molholm et al., 2002)), dovetailing with evidence that multisensory events capture attention and modulate processing in parietal cortex (Bahrick & Lickliter, 2000; Talsma et al., 2007). Critically, although there is not yet consensus on the matter and MSI effects are seen in both subcortical and cortical neurons in anesthetized animals (Allman et al., 2009; Meredith & Stein, 1983), it has been argued that some MSI-related effects are attention-dependent (Talsma et al., 2007; Talsma et al., 2010). More broadly, parietal mechanisms integrate spatial/feature cues to direct attention (Tan et al., 2015) and orchestrate top-down with bottom-up influences (Li et al., 2010), providing a plausible route by which MSI enhances early parietal responses. In contrast, the absence of parietal ERP MSI in AU and SIBs suggests a selective reduction of attentional modulation for multisensory events, rather than a generalized deficit in unisensory encoding. In other words, the NA-specific enhancement appears to reflect attention-mediated multisensory gain, which is absent in the AU and the SIB groups, consistent with an attentional account of altered MSI. This idea is further supported by the results observed concerning α-ERD, which is a marker of task-related cortical excitability. α-ERD facilitates the processing of task-relevant stimuli (Foxe et al., 1998; Foxe & Snyder, 2011; Klimesch, 1999; Klimesch et al., 2007; Worden et al., 2000), and is shaped by both bottom-up stimulus characteristics and top-down cognitive demands (Hanslmayr et al., 2011; Peng et al., 2015). In our data, the MSI effect on α-ERD was significantly reduced in autism. As with the ERP findings, α-ERD differences were specific to the multisensory condition, suggesting that unisensory processing is relatively intact in autism, while the mechanisms supporting multisensory processing are selectively disrupted. Notably, the topographies differed across measures: the ERP MSI effect in NA localized over left centro-parietal sites, whereas the α-ERD MSI effect was quantified over parieto-occipital channels, consistent with modulation of visual cortex excitability during MSI. This suggests that the ERP may index a more precise attentional-parietal mechanism engaged during multisensory processing, whereas the α-ERD effect reflects a more diffuse reduction in cortical excitability modulation in autism.

Finally, diminished theta-band connectivity between frontal and parieto-occipital regions during AV processing points to impaired long-range communication, likely reflecting disrupted top-down attentional control. These effects were not observed during unisensory trials, reinforcing that these differences are specific to multisensory contexts. Our findings extend previous evidence of atypical MSI in autism (Brandwein et al., 2013; Brandwein et al., 2011; Noel et al., 2018; Russo et al., 2010) and underscore the value of integrating ERP, oscillatory dynamics, and connectivity measures to better characterize sensory dysfunction in neurodevelopmental disorders.

### Impaired Functional Connectivity and Top-Down Modulation During Multisensory Processing

In autistic children, theta-band functional connectivity between visual and frontal regions was reduced during multisensory stimulation. This diminished wPLI within 110–280 ms suggests an early breakdown in long-range coordination between sensory and higher-order control networks such as the dorso-lateral prefrontal cortex (DLPFC). The DLPFC is widely implicated in top-down attentional control (Katsuki & Constantinidis, 2012; Kondo et al., 2004; Rossi et al., 2009), and reduced connectivity with occipital regions may reflect a failure to effectively modulate sensory processing in autism (Chen et al., 2024; Just et al., 2012). Automatic attentional capture by multisensory events has been proposed as a key mechanism underlying multisensory gain (Talsma et al., 2010; Van der Burg et al., 2008). Bahrick and Lickliter (Bahrick & Lickliter, 2000) suggest that intersensory redundancy guides attention and perceptual learning during development. If this mechanism is compromised in autism, it could help explain why multisensory benefits are reduced, as suggested by prior work showing atypical MSI effects only under conditions requiring attentional modulation (Magnee et al., 2011). The breakdown of top-down coordination observed here may therefore underlie the reduced cortical activation we report for autistic children, pointing to a broader failure to dynamically integrate multisensory inputs, a process essential for adaptive perception and behavior. Supporting this interpretation, we found that stronger parieto-occipital–DLPFC connectivity was associated with larger P1 and N1 amplitudes in NA and SIB groups, whereas this relationship was absent in AU (Supplementary Figure 6). This dissociation reinforces the view that disrupted connectivity compromises the ability of attentional control systems to enhance early sensory responses in response to multisensory stimulation in autism.

### Behavioral Implications and Neural–Behavioral Relationships

Behaviorally, AU and SIB participants showed reduced multisensory facilitation, evidenced by the absence of race-model violation on repeat trials, indicating less benefit from simultaneous auditory–visual input than NA peers. Although α-ERD did not correlate with our behavioral MSI metrics, within the AU group, α-ERD was significantly associated with reaction time, suggesting that weaker cortical engagement relates to slower responses (e.g., greater processing demands or compensatory strategies). In addition, theta-band wPLI between parieto-occipital regions and DLPFC correlated with reaction time in NA but not in AU, with SIBs showing an intermediate pattern (Supplementary Fig. 6). This dissociation may point to a disruption in the usual coupling between top-down attentional control and performance in autism, consistent with inefficient functional coordination during multisensory processing. Importantly, we did not observe significant correlations between neural MSI markers (ERP or α-ERD AV − A+V) and behavioral MSI (area under the race-model violation curve or the maximum within the first four quartiles). One possibility is that the simplicity of the task limited behavioral MSI, reducing power to detect brain–behavior links. Future work using more demanding, noise-degraded multisensory stimuli may elicit stronger behavioral facilitation and clarify how neural MSI relates to performance.

### Familial Traits: Intermediate Connectivity and Alpha Suppression in Siblings

The current data reveal for the first time that MSI, which many studies, including the current one, have shown to be impaired in children with autism, is also impaired in biologically-related first-degree relatives. This was evident in statistically reduced behavioral indices of MSI in the SIB compared to the NA group. In addition to this behavioral finding, the SIB group exhibited an electrophysiological profile that appeared to fall between the NA and AU groups: for MSI related measures that the AU and NA groups differed statistically on, visual inspection of the data suggested an intermediate pattern in SIBS. However, this intermediate pattern did not lead to statistically significant differences from either the NA or AU group. While these data are consistent with these measures representing inherited vulnerability for autism, further work is needed to test this interpretation and understand the robustness of these effects. Nevertheless, the fact that this intermediate pattern emerges consistently across both early sensory (α-ERD) and top-down connectivity (wPLI) measures supports the idea that these neural features may index underlying biological vulnerability within the broader autism phenotype, even in the absence of overt behavioral symptoms. This interpretation aligns with prior findings showing that siblings of autistic children can exhibit subclinical or partial expressions of autism-related neurophysiological traits (Matthews et al., 2013; Narvekar et al., 2022).

### Importance of Eye-Tracking Control

A critical methodological aspect of this study was the use of eye-tracking data to exclude trials in which participants were not looking at the center of the screen. Without this gaze-based filtration, group differences in P1 amplitude also emerged in the visual-only condition, likely due to increased variability in gaze position among autistic children. Given that early visual components such as P1 are highly sensitive to retinal stimulus location (Frey et al., 2013; Kim et al., 2016), failing to control for fixation can introduce confounds and lead to inaccurate interpretations of group differences. These findings underscore the importance of incorporating eye-tracking measures in neurophysiological studies, particularly for studies that include visual stimulation in clinical populations where gaze behavior may be atypical.

### Limitations

This study has several limitations. First, although we applied a surface Laplacian transform to improve spatial specificity, interpreting EEG-based connectivity remains challenging due to the inherently low spatial resolution of scalp recordings (Haufe et al., 2013). While our findings indicate reduced theta-band connectivity between sensory and frontal regions during MSI in autism, confirming the precise spatial origins and directionality of these effects will require complementary modalities such as fMRI and intracranial EEG, which offer higher spatial precision for the characterization of large-scale network dynamics.

Second, our sample consisted of autistic children without intellectual impairment and with functional language, representing only a subset of the autism spectrum. These findings may not generalize to individuals with lower cognitive or language abilities. Finally, although we restricted the age range to reduce variability, multisensory integration develops differently in autistic and non-autistic groups. Longitudinal or cross-sectional studies examining developmental trajectories would provide important insights into how MSI and its neural correlates evolve over childhood (Crosse et al., 2022; Cuppini et al., 2017; Foxe et al., 2015; Monti et al., 2023).

### Conclusion and Future Directions

Taken together, our results demonstrate that autistic children exhibit reduced cortical activation and diminished long-range connectivity during multisensory processing. These deficits were specific to the audiovisual condition, reinforcing the idea that multisensory processing is particularly disrupted in autism. Together, significantly attenuated MSI effects on ERPs and α-ERD, and decreased theta-band wPLI between parieto-occipital and frontal regions suggests that atypical MSI in autism reflects combined impairments in multisensory modulation of local cortical excitability and large-scale network communication. Importantly, unaffected siblings displayed an intermediate neural profile on both α-ERD and functional connectivity measures, showing no significant difference from either NA or AU groups. Although requiring further study with larger sample sizes, this pattern is consistent with the possibility that these neural alterations reflect familial or endophenotypic traits associated with the broader autism phenotype, even in the absence of clinical symptoms. Future research should examine whether interventions targeting neural excitability, connectivity, and attentional regulation (such as neurofeedback, rhythmic entrainment, or non-invasive brain stimulation) can improve multisensory integration in autism. Moreover, identifying early electrophysiological markers of multisensory dysfunction, including those observables in unaffected siblings, may aid in stratifying individuals on the spectrum and tailoring intervention strategies during development.

## Author contributions

S.M, J.J.F conceived the study. S.B contributed to experimental implementation. T.V preprocessed and analyzed the data under the supervision of S.M. T.V wrote the first draft of the manuscript. S.M., S.B., and J.J.F provided edits on the manuscript. All authors approved the final manuscript.

## Acknowledgment

We thank Dennis Cregin, Trinca Lecaj, Daniella Coen, and Albulena Sejdu for their essential contributions to participant recruitment and data acquisition. This work was supported by a grant from the Simons Foundation Autism Research Initiative (SFARI Award # 874845, SM). Support for recruitment and phenotyping of participants was provided by the Human Clinical Phenotyping Core of the NICHD funded Rose. F. Kennedy Intellectual and Developmental Disabilities Research Center (P50 HD105352, SM). Work at the collaborating site in Rochester is supported through the Golisano Intellectual and Developmental Disabilities Research Institute (UR-IDDRC), which is supported by a center grant from the Eunice Kennedy Shriver National Institute of Child Health and Human Development (P50 HD103536, JJF). The content is solely the responsibility of the authors and does not necessarily represent the official views of the National Institutes of Health.

## Footnotes

1 In this paper, we primarily adopt identity-first language, reflecting prevailing preferences among many autistic adults and self-advocates. We recognize, however, that person-first language is preferred by others in the autism community.

---

**Supplementary figure 1.**
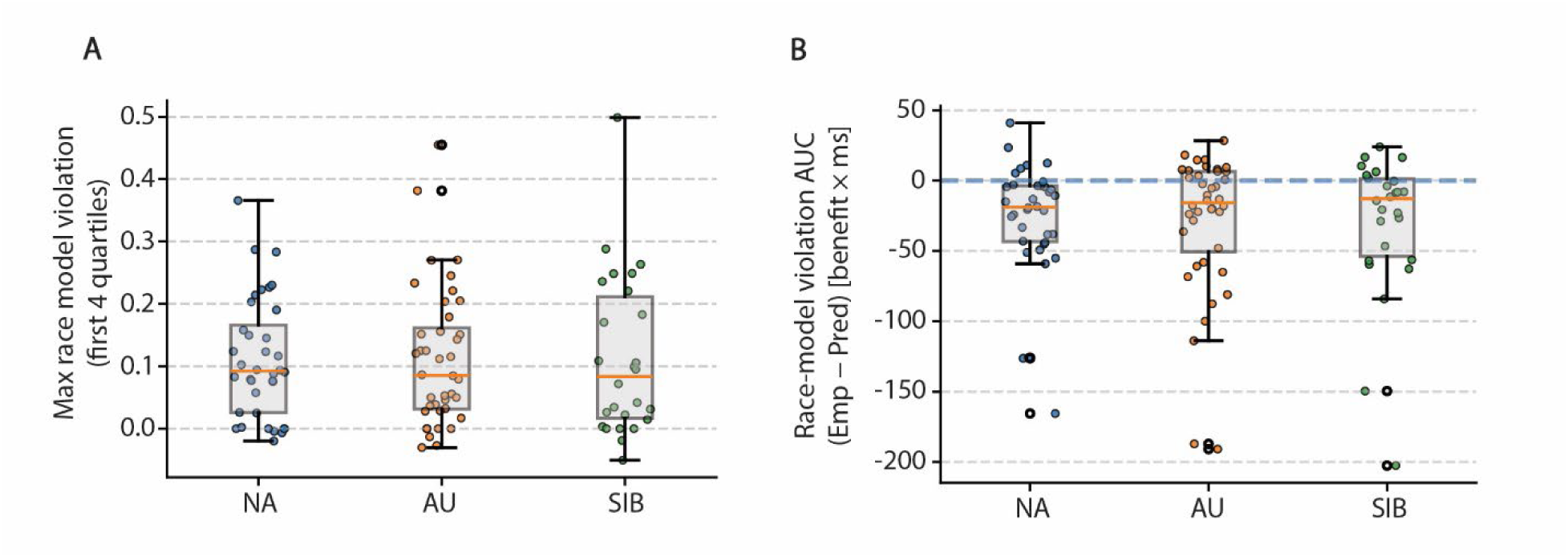
Absence of statistical differences of race model violation between groups.

**Supplementary figure 2.**
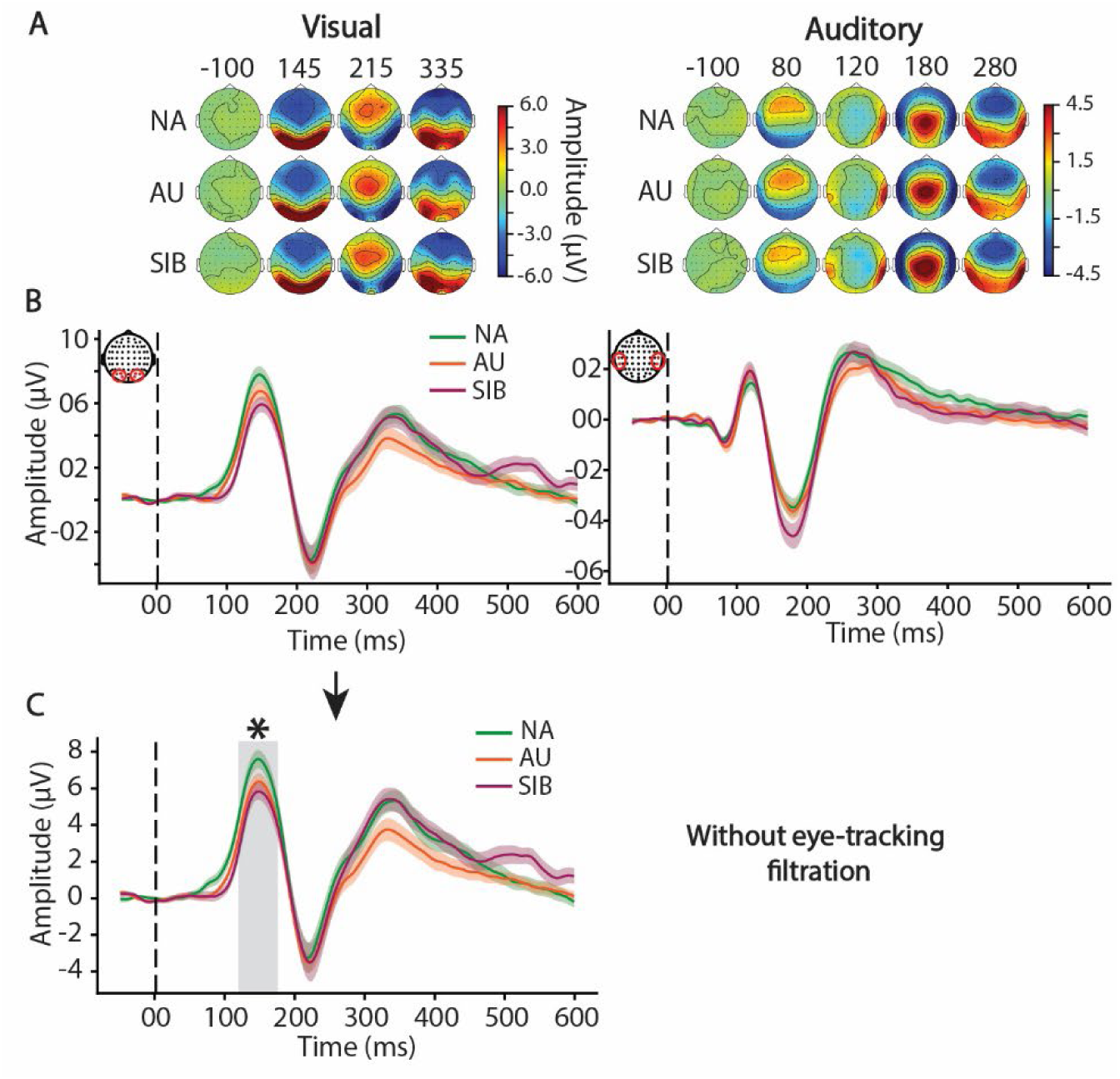
**Effect of eye-tracking filtration on ERPs.**

**Supplementary figure 3.**
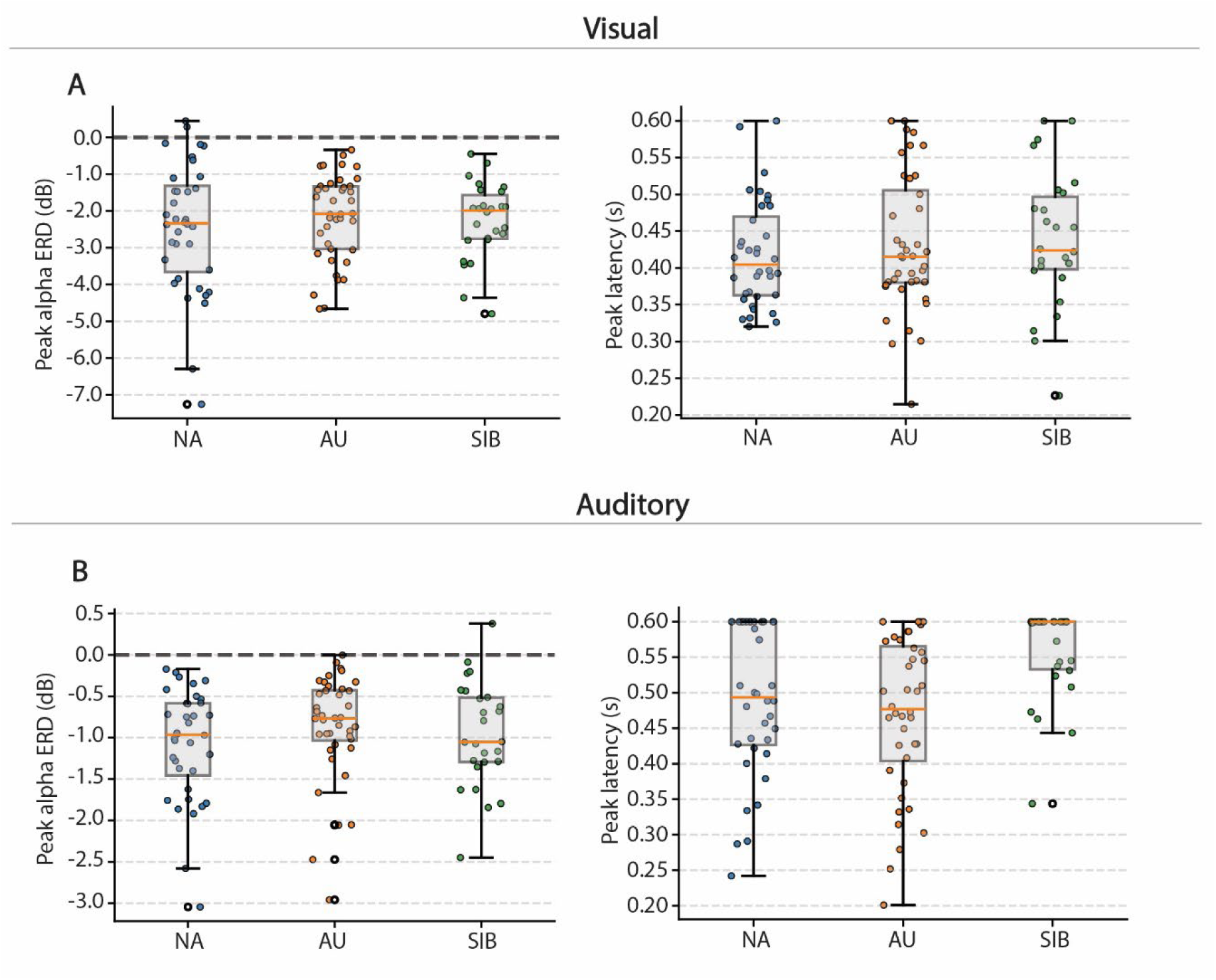
**α-ERD Peak amplitude and latency for unisensory stimulation.**

**Supplementary figure 4.**
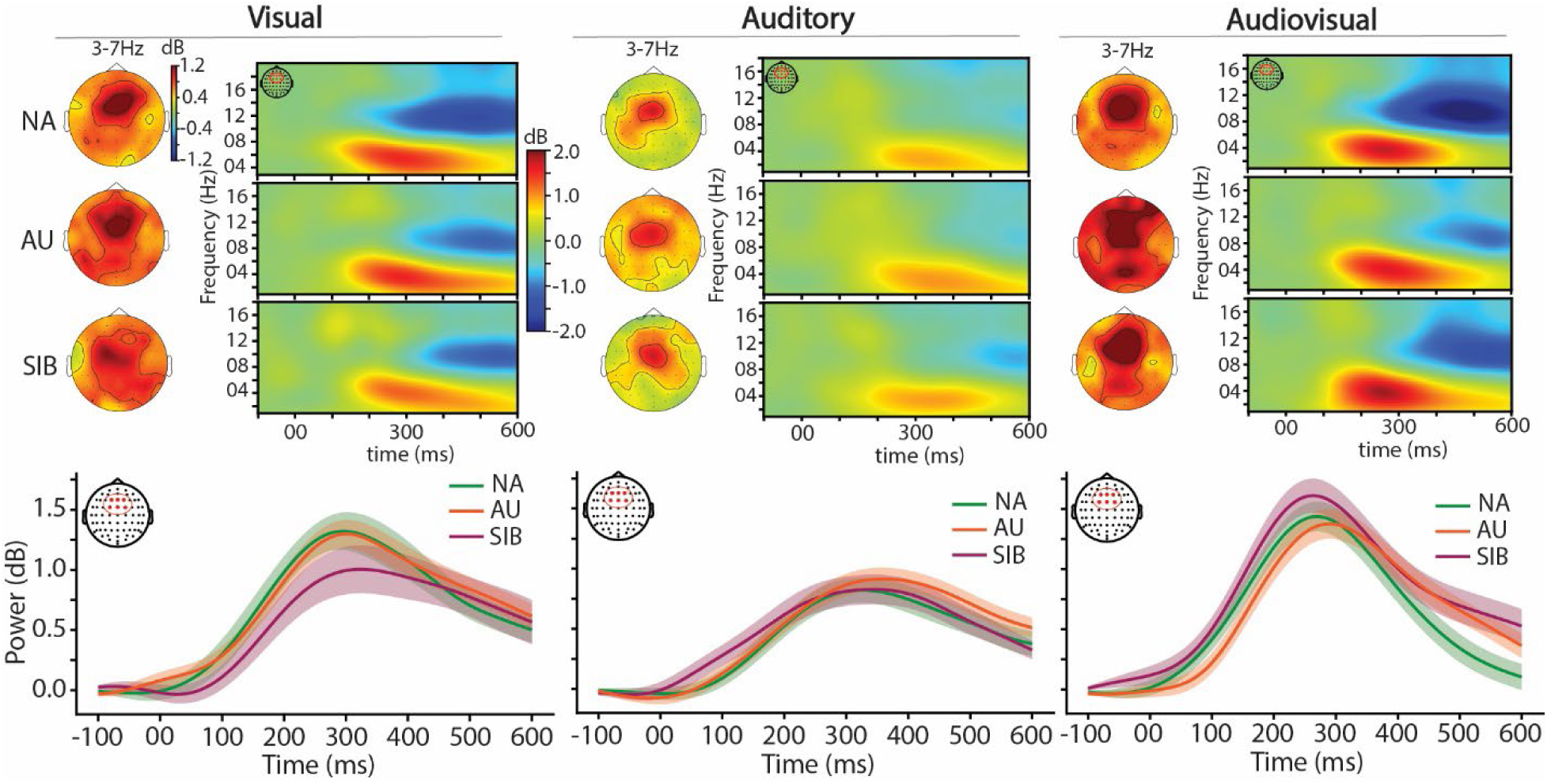
Comparable Increases in Induced Theta-Band Activity Across Groups in Response to Different Sensory Modalities. (**A**) Averaged topographical distribution of induced theta activity (4–7 Hz) in response to visual (left), auditory (center) and audiovisual (right) stimulation along with the corresponding time-frequency map for a cluster of occipital channels in response to the visual stimulation for NA (top row), AU (middle row), and SIB (bottom row). (**B**) Induced theta activity averaged over a cluster of centro-frontal channels (see corresponding topomaps) for NA (green), AU (orange), and SIB (purple) in response to the visual (left), auditory (center) and audiovisual (right) stimulation The shaded areas in the plots represent the standard error of the mean (SEM) in the corresponding colors.

**Supplementary figure 5.**
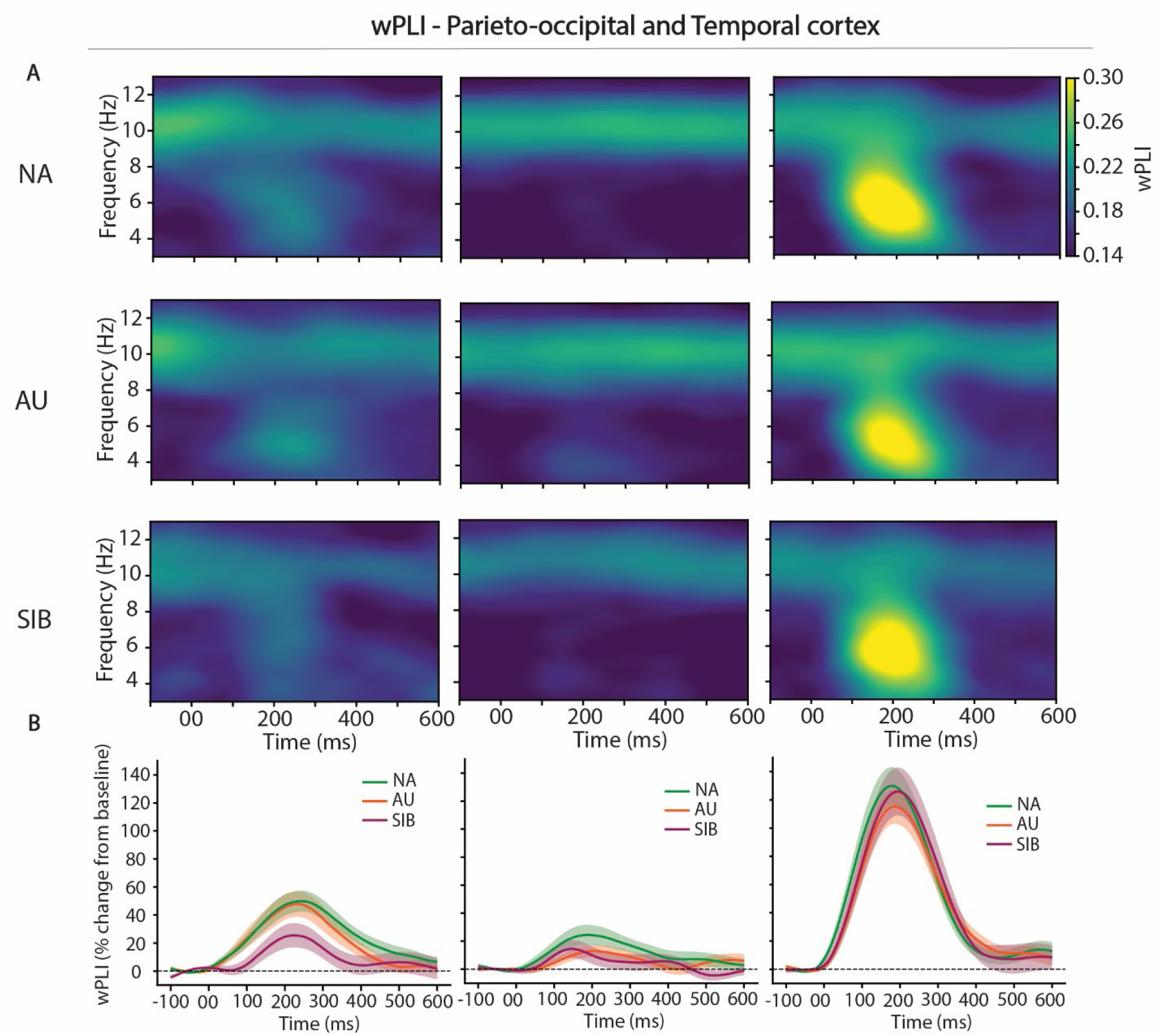
wPLI between parieto-occipital and temporal cortex. (**A**) Time-frequency representation of the weighted-phase lag index (wPLI) between a cluster of parieto-occipital channels and a cluster of temporal channels in response to a visual stimulation (left), auditory stimulation (center) and audiovisual (right) for the NA group (top row), AU group (middle line) and SIB group (bottom line). (**B**) wPLI averaged for the theta band (4-7Hz) for NA (green), AU (orange), and SIB (purple) for the corresponding sensory stimuli.

**Supplementary figure 6.**
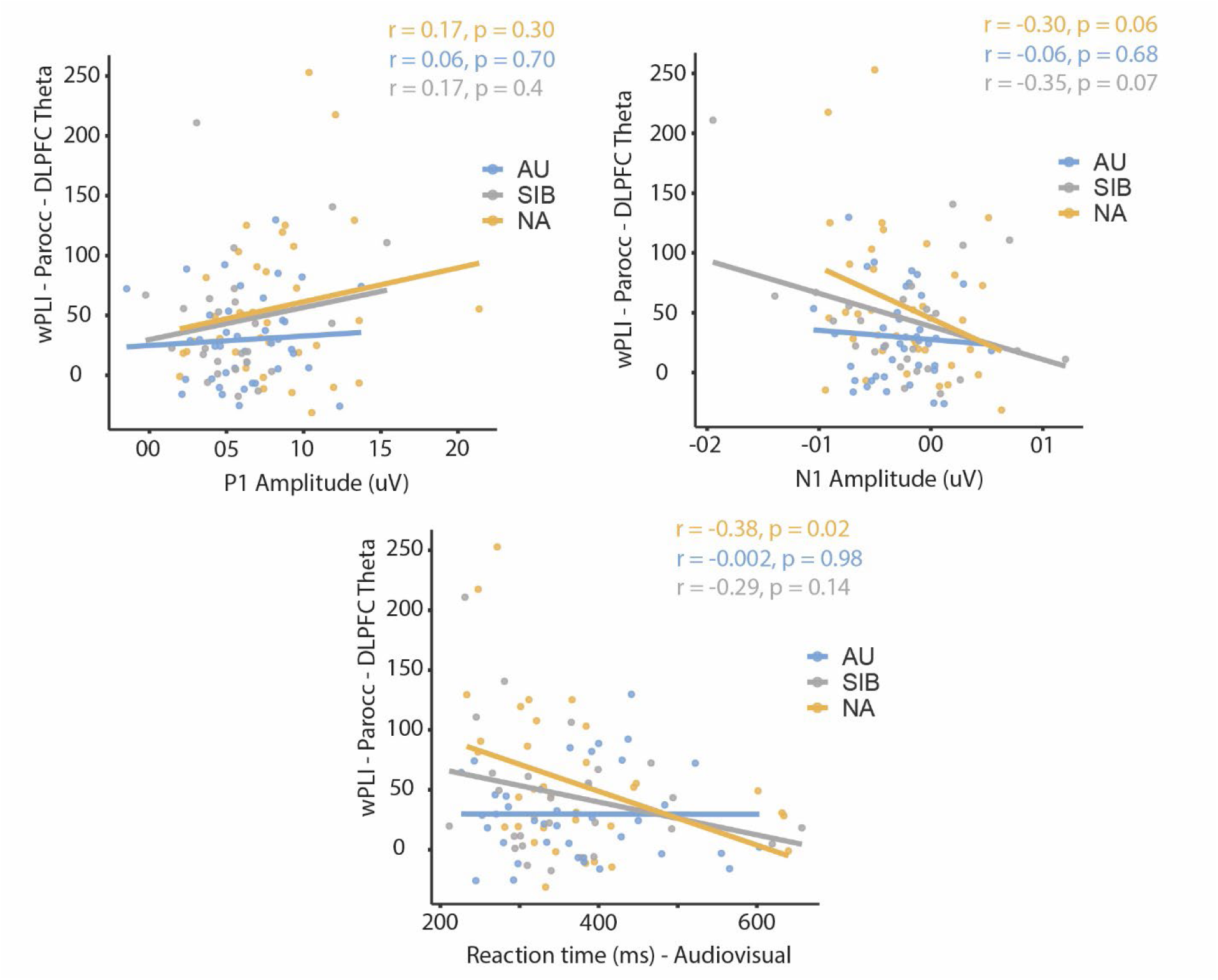
Correlation between wPLI and ERPs components and Reaction times.

